# No evidence for disassortative mating based on HLA in a small-scale, endogamous population

**DOI:** 10.1101/2025.05.06.652536

**Authors:** Gillian L. Meeks, Brooke Scelza, Katherine. M. Kichula, Catrinel Berevoescu, Kristin Hardy, Ticiana D.J. Farias, Genelle F. Harrison, Nicholas R. Pollock, Neus Font-Porterias, Sean Prall, Paul J. Norman, Brenna M. Henn

**Author notes:** Indicates co-authorship position (Gillian L. Meeks, Brooke Scelza, Paul J. Norman, Brenna M. Henn).

## Abstract

Studies dating back several decades have suggested that humans prefer potential mates with dissimilar HLA genotypes. Evidence for actualized disassortative mating based on the human-specific MHC remains inconclusive. For instance, cosmopolitan populations have often exhibited the opposite trend whereby assortative mating at the MHC is observed, indicating that social stratification may overwhelm potential biological mate preferences. However, small-scale, endogamous populations–whose social structures more closely resemble those throughout most of human evolution–have been largely overlooked. Here, we assess HLA dissimilarity among Himba pastoralists from Namibia, where socially accepted concurrency allows individuals to maintain both arranged marital and self-selected (“love match”) partnerships. This provides a rare opportunity to directly test HLA similarity across contrasting partnership types (arranged vs chosen) within the same social system (n = 249 observed partnerships). We find no difference in HLA dissimilarity (neither at the genotype nor protein divergence level) between partnership types, nor in their fitness benefits to potential offspring as assessed via computationally predicted pathogen binding affinities. The effects of the partnership types likewise do not differ from a random, background distribution of 18,487 possible unrelated pairings. Finally, we detect extensive haplotype sharing across the HLA region, suggesting that episodes of fluctuating positive selection may be a stronger force maintaining HLA polymorphism than disassortative mating, even in an evolutionarily relevant social context.

## 1. Introduction

MHC dissimilarity is predicted to play a critical role in mate choice strategies, as choosing a dissimilar mate enhances offspring immune function by increasing their heterozygosity and therefore broadening pathogen resistance. Individuals who are heterozygous are codominant for MHC allele expression. Sexually mediated balancing selection has therefore been proposed as a major force maintaining extreme allelic diversity in this region (1–3). Meta-analysis of 27 non-human vertebrate species has indeed shown a small, but significant preference for MHC dissimilar mates (4). Earlier experimental work at the human specific MHC (i.e., HLA) demonstrated both male and female preference for the odour of HLA dissimilar mates (5). Initial studies about realized mate choice in two ethnically and genetically homogeneous populations (Hutterites and Mormons) likewise reported greater HLA dissimilarity among married partners than expected under random mating (6, 7). However, meta-analyses of the cumulative literature in the decades since have found no significant mate preferences for HLA dissimilarity (8, 9). In fact, when ethnically heterogeneous, large-scale study populations are considered, the opposite pattern emerges: assortative mating is more common, where partners are more similar at their HLA than random pairings within the population (8). Cosmopolitan populations in the Western Hemisphere tend to reflect migration from multiple continental and ancestral sources and in such populations, married pairs often share more similar ancestry proportions than random pairings (10–12). Such ancestry-assortative mating can arise either from active preference for partners of similar ancestry proportions within an admixed population, or from socially-structured mating between groups that contain ethnic, linguistic, or socioeconomic diversity (13).

One way to test this idea is to directly distinguish between HLA and overall genome-wide relatedness. In comparisons across six European and one Israeli population, both the Dutch sample and a joint analysis of five Northern European populations did show significantly greater HLA dissimilarity among married pairs than in randomly permuted pairs, independent of genome-wide similarity (14). These results suggest that preference for HLA-dissimilar mates can sometimes be detected in cosmopolitan populations with sufficiently large sample sizes. However, another recent meta-analysis of studies that successfully controlled for genome-wide relatedness (including the seven estimates from the aforementioned study) still found no overall evidence for HLA-based disassortative mating (9).

Notably lacking in studies that control for genome-wide similarity are any non-European, small-scale societies. A more evolutionarily relevant test of HLA-based mate preferences would therefore examine smaller-scale, endogamous societies–social structures that more closely reflect those in which human mating behaviors evolved and where ancestry-assortative mating is less likely. Moreover, in heterogeneous cosmopolitan populations, mating with almost any unrelated individual may already yield offspring with dissimilar HLA haplotypes beyond a threshold that maximizes fitness, potentially obscuring subtle selection for HLA dissimilarity. In contrast, in small, endogamous populations where background relatedness is higher, evolved preferences for HLA-dissimilar mates may be more pronounced (9, 15).

We directly test for evidence of mate selection based on HLA dissimilarity, independent of genome-wide similarity, in a group of Himba agropastoralists in northern Namibia–the first study of its kind in a non-European, small-scale society, and with a substantially larger sample than the only previously analyzed small-scale population controlling for genome-wide similarity (HapMap CEU individuals from the Mormon community n=28 (7), n=24 (15)). Several features of the Himba marital system make them particularly well-suited for this investigation. Marriages may be either arranged (via parental choice) or self-selected (“love matches”) (16)) and concurrent partnerships are both common and socially acceptable (17–19). If intrinsic preferences for HLA-dissimilar mates exist, we would expect to observe them among love matches rather than arranged marriages. Moreover, arranged marriages often fulfill social obligations, potentially allowing love matches to reflect more active mate choice (17–19)). The Himba are also genetically well-suited for studying ancestral mate preferences. While they are endogamous, they exhibit greater overall genetic diversity than the previously studied out-of-Africa isolates, like the Hutterites and Mormons, making them more relevant for inferences about the majority of human evolution. Finally, admixture from neighboring populations is minimal (mean Western African ancestry >95% across k = 4,5 runs), and the Himba have served as a reference “Western African” population in our group’s admixture analyses (20), suggesting that ancestry-assortative mating is unlikely to confound results.

We use paired genetic HLA and genome-wide genetic data along with partnership records to test whether unions resulting from partner choice exhibit greater HLA dissimilarity–at both the genotype and protein sequence level–than arranged marriages. Additionally, we assess the predicted pathogen-binding capacity of hypothetical offspring from arranged and chosen partnerships to evaluate potential fitness benefits of parental HLA allele combinations.

## 2. Methods

### (a) Study Population

The Himba are an agro-pastoralist group residing in Northwestern Namibia and Southern Angola, who have a total population size of around 30,000-40,000 individuals (21). The groups remain semi-nomadic but increasingly have access to formal education and the market economy. The present study was conducted in the community of Omuhonga as part of the Kunene Rural Health and Demography Project, which has been working in the community since 2010. There are about 1000 residents of Omuhonga, living in extended family compounds. Marriages are arranged by parents or other kin, although “love match” marriages are common, particularly after the first marriage (16). Extra-marital relationships are also common for both men and women (17).

### (b) Data Generation

DNA was collected with informed consent from individuals via saliva using the prepIT-L2P kit and protocol. DNA was used to generate HLA genotype calls for 366 individuals. HLA class I and II loci were targeted for DNA sequencing using a well-established biotinylated DNA probe-based capture method (22). HLA alleles were determined from the sequence capture using the consensus calls obtained from two algorithms: NGSengine® 2.10.0 (GenDX, Utrecht, the Netherlands) and HLA Explore™ (Omixon Biocomputing Ltd. Budapest, Hungary). Alleles were then converted into their P-group designations from https://hla.alleles.org as of 10/7/2024 (2-field designation), which groups alleles whose nucleotide sequences encode identical protein sequence in the peptide binding domains (exon 2 and 3 for HLA class I and exon 2 for HLA class II alleles). Genotype array data using one of two arrays, Illumina H3Africa or MEGAex, were previously generated for 360 of these individuals [dbGaP: phs001995.v3.p1]. One individual was newly genotyped using H3Africa. Five individuals with HLA capture data did not have corresponding SNP array data and were not included in the partnership analyses requiring genome-wide relatedness metrics. These SNP array datasets previously underwent quality control measures and were merged as previously described (18).

There were 219 adults (82 males, 137 females) having both genotype array and HLA genotype data who were in at least one documented partnership: arranged marriage, love match marriage, self-reported informal partnership, or a discovered informal partnership. Marriages and self-reported informal partnerships were recorded during demographic interviews conducted between 2010 and 2017. Additional informal partnerships were discovered due to shared biological offspring as determined by a list of parent offspring trios generated via KING 2.1.3 (23), using the identical by descent (IBD) segment flag. There were 47 arranged marriages, 47 love match marriage pairs, and 155 informal pairs. Of the informal pairs, 48 were discovered only through shared biological offspring. These pairings may have been excluded during the interviews because they occurred many years in the past and our interviews only asked about current partnerships. The 47 love match marriages were combined with 203 informal partnerships and designated as chosen partnerships to compare with the 47 arranged partnerships in our analyses.

### (c) Genotype array data quality control

Individuals were genotyped using one of two arrays, H3Africa or MEGAex. These datasets previously underwent quality control measures and were filtered using PLINK v1.90b3v(24) for missingness greater than 5%, a MAF less than or equal to 1%, and a Hardy–Weinberg equilibrium exact test with a *p*-value below 0.0001 as described previously (18) and documented in dbGaP accession phs001995.v3.p1. The merged array dataset containing 376,657 SNPs was used for analyses.

### (d) Creating a set unrelated sets of individuals

Traditional pedigree inference methods that use expected IBD sharing perform poorly in endogamous populations with elevated background sharing and pedigree reticulations. We used PONDEROSA (25), recently developed in our lab, to accurately infer close relationships in our sample. PONDEROSA uses a machine learning approach to learn the population distribution of IBD summary statistics to more accurately classify close relationships. Based on the inferred relations we excluded any pairs who were 2^nd^ degree or closer relatives from our background distributions in our linear models (Statistical Methods eq 2-3). Third degree relative pairs were permitted in this background distribution to reflect the stated cultural preference for cousin marriages in arranged unions (16). We also used this program to maximize a set of unrelated individuals related at 4th degree or less (n=102 individuals) for allele and haplotype frequency based analyses.

### (e) Hardy-Weinberg Analysis

To test whether the HLA genotypes complied with Hardy-Weinberg equilibrium, we used the Asymptotic Statistical Test with Ambiguity (ASTA) from the HWETests package (26). ASTA is designed to evaluate deviations from Hardy-Weinberg equilibrium for highly polymorphic loci while accounting for ambiguity in genotype calls. Using ASTA, we ran the full_algorithm() function with default parameters on the set of 102 unrelated individuals (maximum relation of 4th degree).

### (f) Identity by Descent (IBD) Sharing

Genotype data was phased using SHAPEIT v2.r837(27) with --duohmm -W 5 using an ancestry-matched recombination map (28), constructed based on linkage disequilibrium patterns in the 1000 Genomes Project (29) Yorubans, matching the predominant West African ancestry present in the Himba. We called IBD segments with phasedIBD (30) using default parameters with phase correction turned off. We determined phasedIBD calls to be slightly more accurate than germline 1.5.3 (31) and GERMLINE2 (32) by comparing the total cM of IBD shared between parent offspring pairs (Table S1). We expected parents and offspring to share the full length of the recombination map in IBD1 and share an additional background amount of IBD2 estimated to be the average cM of IBD shared between unrelated (> 3rd degree relations) individuals in the population. The phasedIBD calls without phase correction most closely matched this expectation. The phasedIBD calls with phase correction option turned on were the least accurate. A minimum threshold of 3cM was used for all runs.

### (g) HLA similarity score

We assessed HLA similarity in terms of the number of alleles shared across the eight typed HLA genes for each pair (minimum of 0 if no alleles are shared at any gene, maximum of 16 if both alleles at each gene are shared). A shared allele in a homozygous state in one individual only counted as one shared allele between the individuals. Importantly, due to the many alleles at each HLA gene, two individuals with heterozygous genotypes at a locus do not necessarily imply shared alleles.

### (h) Average log odds of homozygous offspring

Odds of homozygous offspring for each locus were determined according to standard mendelian transmission probabilities, accounting for the multi-allelic states of each gene. Table S2 shows the possible states for this variable considering only one gene. Importantly, for a given number of alleles shared at a locus, there are multiple possibilities for the odds of homozygous offspring depending on the genotype states of the individuals in the pair. We log transformed these odds to normalize the distribution and then averaged across the 8 loci. We included a +0.1 boundary correction in the numerator and denominator of the odds calculations to allow for numerators and denominators of 0 (eq 1).

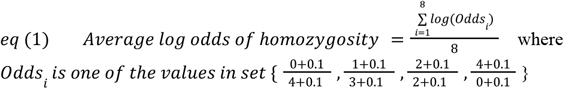

### (i) Peptide Binding Groove Protein Divergence

We used a script available from a previous publication (33) to assess the mean Grantham amino acid distance (34) of the peptide binding groove between HLA molecule allotypes, comparing the 4 pairwise combinations of allotypes between two individuals. Only peptide binding groove sequences (exons 2 and 3 for HLA class I and exon 2 for HLA class II) were compared because all alleles were grouped according to their P group (identical peptide binding grooves). We used the fasta files provided for the HLA class I, DRB1, and DQB1 alleles and the Grantham distance matrix available in a previously published package (https://granthamdist.sourceforge.io/). For the remaining HLA class II alleles we used the IPD-MHC (https://www.ebi.ac.uk/ipd/mhc/alignment) multiple sequence alignment tool to align alleles and extract exon 2 amino acid sequences.

### (j) Pathogen Proteins

We modified a previously published list of representative human pathogen proteins (33) that was curated from the Gideon database (35) based on global distributions, potential for high mortality, and historical relevance. After personal correspondence with an infectious disease expert, Dr. Ashley Hazel, who has extensive experience working with the Himba community we removed 6 pathogens unlikely to be ecologically relevant in Northern Namibia: *Entamoeba histolytica* (amebiasis), *Trichinella spiralis* (roundworm), *Schitosoma mansoni* (schistosomiasis), HIV, Hepatitis C, and *Yersinia pestis* (plague). We added 4 sexually transmitted diseases known to be highly prevalent in the Himba (HSV1, HSV2, *Chlamydia trachomatis* (chlamydia), and *Neisseria gonorrhoeae* (gonorrhea)) for a total of 25 pathogens (36). We used the NCBI accession numbers available in the previously published study (33) to gather amino acid sequences for the antigenic proteins of the 21 pathogens overlapping that publication. For the 4 additional pathogens we used IEDB https://www.iedb.org/ to find the pathogen proteins that are MHC ligands and then identified these proteins’ amino acid sequences using NCBI. The complete list of 298 proteins and their accession numbers are available in Appendix 1.

### (k) Pathogen Binding Prediction

We used NetMHCpan (v4.1) (37) to predict peptide binding affinity for the allotypes of HLA class I (HLA-A, B, and C) and used NetMHCIIpan (v4.3) (38) for the allotypes of HLA class II (HLA-DP, DQ, and DR). These algorithms are trained using experimental binding affinity and eluted ligand data to predict binding affinity. Binding prediction was performed on all possible amino acid 9mers to match the 9-residue binding groove for HLA class I genes and all possible amino acid 15mers for HLA class II molecules. The HLA class II molecules DPα1 and DPβ1, DQα1 and DQβ1, and DRα1 and DRβ1 can often combine to form the DP, DQ, or DR epitopes, respectively (39, 40). DRα1 has limited polymorphism, and we did not include DRβ3/4/5 in this analysis so only DRβ1 binding was assessed for DR. All possible combined DP and DQ molecules were cross-referenced for their binding affinities. The predicted binding affinity scores between every possible peptide and every MHC molecule were ranked as a percentage by the algorithms based on comparison with a large set of naturally occurring peptides. We used the default rank threshold for strong binding peptides (top 0.5% for HLA class I genes, and top 1% for HLA class II genes). One of the rare alleles (DPB1*61:01N) is a null allele, i.e., does not present peptides, and thus pairs including the individual with this allele were excluded for the DP binding analysis.

### (l) HLA haplotype resolution

The HLA allele calls from target sequence capture are unphased. We resolved HLA haplotypes by imputing closely matched HLA allele calls from the paired, phased SNP data across the 5Mb region encompassing the typed HLA genes. Although HLA imputation based on sparse genotype data is not accurate at detailed resolutions, we were still able to resolve the haplotypes by comparing imputed alleles to the known alleles from the HLA targeted sequencing. We trained HIBAG HLA imputation classifier models (41) on the Himba SNP data across the whole 5Mb region with paired HLA allele calls from the target capture (n=361 individuals). The models had “out of bag” accuracies of 97.96%, 96.97%, 98.60%, 98.75%, 98.73%, 98.75%, 97.66% across A, B, C, DPB1, DQA1, DQB1, and DRB1, respectively. The program does not impute DPA1 genotypes. We then extracted each phased SNP-based haplotype at the HLA region and created two pseudo-homozygous genotypes for each individual (i.e., ID1_hap1|ID1_hap1 and ID1_hap2|ID1_hap2). We ran these pseudo-homozygous genotypes through our curated HIBAG model to predict the HLA allele associated with each pseudo-homozygous genotype. We ensured that the predicted alleles across the two pseudo-homozygous genotypes matched one of the known target sequenced HLA allele calls for at least the first field. In this way, we filled in the allele calls across the two phased haplotypes. We were able to resolve 698/722 haplotypes with this method. The linkage disequilibrium patterns determined via asymmetric conditional linkage disequilibrium closely correspond to previously identified patterns (42) with high LD between HLA-B and C and also among HLA-DRB1, DQA1, and DQB1, demonstrating the accuracy of our haplotype resolution (Figure S1).

### (m) Calculating IBD sharing rates

To calculate IBD sharing rates at each base pair genome-wide, we used a line sweep algorithm (Figure S2). In this method, the start of an IBD segment is marked +1, and the base pair directly following the last base pair in the IBD segment, is marked -1. A ‘line’ sweeps across the chromosome-length array summing the tally at the current base pair with the tally carried over from the previous base pair to generate IBD coverage scores for each base pair. In this way, a base pair located within a larger set of IBD fragment markings receives a larger coverage value during the line sweep step. The IBD sharing rate for each base pair is then computed by dividing the IBD coverage score by the total number of possible haplotypes in the dataset, the number of pairs multiplied by 2. This procedure reduces the complexity of the IBD sharing calculation problem, allowing for efficient computation of the IBD sharing rate at each base pair, genome-wide. IBD sharing rates were binned and averaged in 1000 base pair chunks.

### (n) Statistical Models

We constructed a dyadic data model (43) to assess the effect of partnership type on IBD sharing (eq 2) and on our HLA metrics (eq 3): HLA similarity score, mean log odds of homozygous offspring, peptide binding groove divergence, and mean number of predicted pathogen peptides bound by hypothetical offspring genotypes. Each partnership type (arranged, chosen) was encoded as a binary variable, and random effects were included for each female and each male to account for individual-level variation. IBD sharing was included as a covariate in equation 3 to determine whether differences in HLA metrics between partnership types remained significant after conditioning on overall genetic relatedness.

The intercepts of both models represent a baseline derived from all 18,487 possible opposite sex pairings that are not in known partnerships, are no closer than 3^rd^ degree relatives, and also constrained to those that have a permissible age differential based on the range observed in the known partnership data. This baseline provides a background distribution for the expected similarity of partners in the population if there is random mating, but with avoidance of close kin. We allowed 3rd degree relatives to exist in our background distribution of partners as Himba individuals report a cultural preference for cousin marriages in arranged unions (16). However, surprisingly, no individuals in actual arranged partnerships were related at 3rd degree or closer, but 1 “love” marriage and 2 informal partnerships discovered through pedigree analysis were found to be 3rd degree relatives.

This background distribution allows us to test whether each pair type deviates from the baseline expectation of similarity while providing additional information to estimate the random effects. The random effects of each female and male capture the unique contributions of each individual to their dyadic HLA metrics/IBD sharing across all pairings (both real and random) (43). This approach allows us to identify whether individuals’ real partnerships (arranged and chosen) differ systematically in their similarity scores when compared to their average similarity with potential random partners in the population.

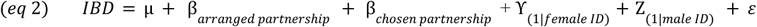

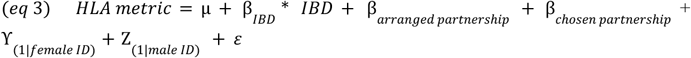

Pairwise contrasts in the effect size estimates of the two different partnership types (arranged vs chosen) were assessed by calculating the Z-statistic, defined as the contrast estimate divided by its standard error. P-values for these Z-statistics were derived from a two-tailed normal distribution. Statistical significance for the main effects and pairwise contrasts was determined using a p-value threshold of 0.05.

## 3. Results

To help contextualize our findings, we first compared HLA diversity in the Himba, which has not been previously characterized, with other publicly available populations, curated to represent a diverse world-wide sample (44) and all available sub-Saharan African groups (45). With HLA allele calls from 366 individuals, we assessed the diversity of HLA alleles based on their P group designation, that is, alleles were grouped according to identical peptide binding domains to indicate differences in pathogen binding function between alleles, across three HLA class I (A, B, C) and five HLA class II genes (DPA1, DPB1, DQA1, DQB1, DRB1) (Figure S3). In order to make unbiased comparisons of allelic diversity based on sample size, we subset our sample to the maximum set of unrelated Himba individuals (n=102, maximum 4th degree relations) (Figure S4). Despite endogamy and a recent population bottleneck (46), the allelic diversity at each gene falls along the linear prediction based on sample size constructed from the publicly available data, with expected or somewhat greater diversity than expected given the Himba sample size (Figure 1). We ensured genotype frequencies at each HLA gene did not deviate from Hardy Weinberg equilibrium (Table S3) as this could indicate genotyping error or population substructure (47).

**Figure 1.**
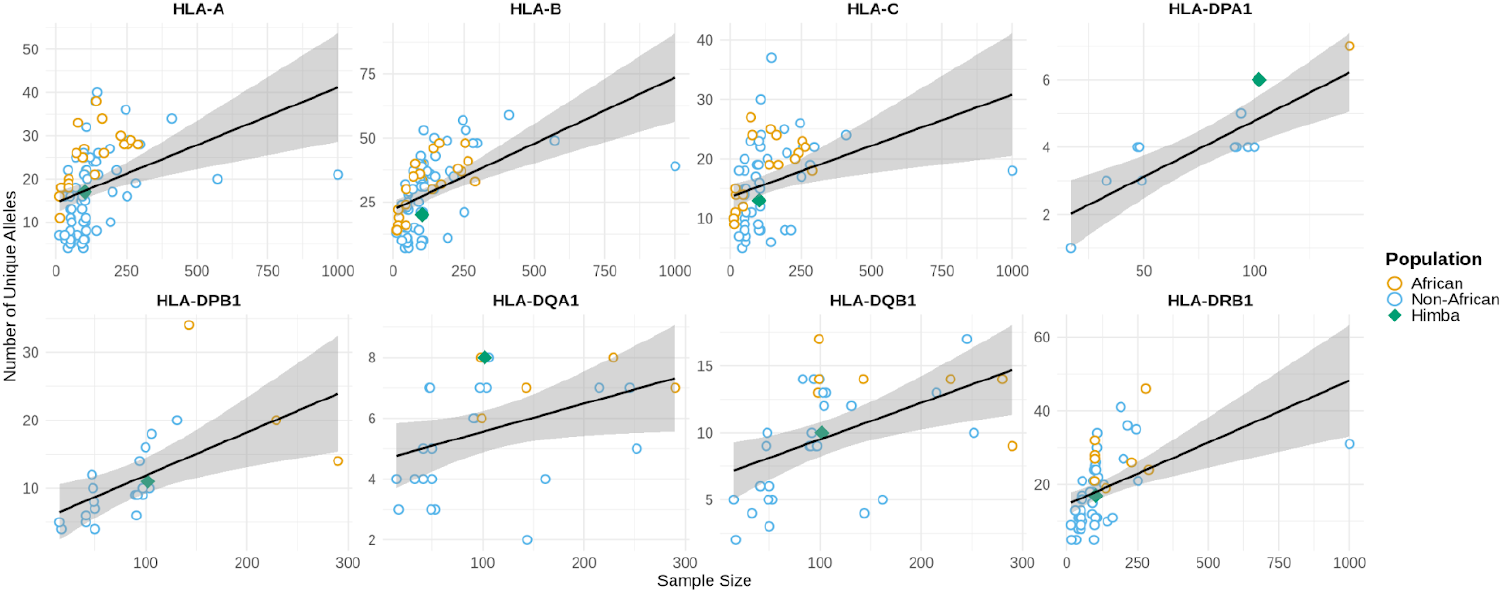
Allelic diversity in the Himba follows or somewhat exceeds linear predictions based on sample size from publicly available cohorts. Allelic diversity for the 8 genotyped HLA genes was measured in the set of 102 unrelated Himba. African cohorts are labeled in orange, non-African cohorts in blue, and the Himba with a green diamond. Linear predictions of the number of unique alleles based on sample size were constructed from the available datasets, excluding the Himba. Ribbons indicate the 95% confidence interval for the linear prediction of the number of unique alleles based on sample size.

Next, we assessed whether there was an effect of partnership type (chosen or arranged) on overall genetic relatedness. If mate choice is, in part, driven by minimizing overall genetic relatedness to reduce fitness consequences associated with increased homozygosity in offspring (46, 48, 49), chosen partnerships could be more genetically dissimilar overall than arranged partnerships. This could produce an associated, but spurious pattern of HLA dissimilarity in chosen partnerships as compared to arranged partners and the background distribution. Separately, preferences for cross-cousin marriage and strong ties between allied families could lead to greater sharing of IBD segments among arranged marriage partners than a background distribution would predict, leading to associated, but spurious patterns of greater HLA similarity in arranged partners as compared to chosen partners and the background distribution.

We assessed overall relatedness via the total number of centimorgans shared in genome-wide IBD segments between the individuals in the partnership (see eq 2 in *Methods*). There was no effect of arranged partnership or chosen partnership on genome-wide IBD sharing, relative to a background population distribution created from random, minimally related (maximum 3rd degree relations), heterosexual pairings with realistic age gaps (Figure 2A) (*p* = 0.84, 0.82, for arranged and chosen partnership effects, respectively) nor a significant difference between the effects of arranged and chosen partnerships in a pairwise contrast of effect size estimates (*p* = 0.93). 3rd degree relatives were allowed in the background distribution to reflect the stated cultural preference for cousin marriages in arranged unions (16).

**Figure 2.**
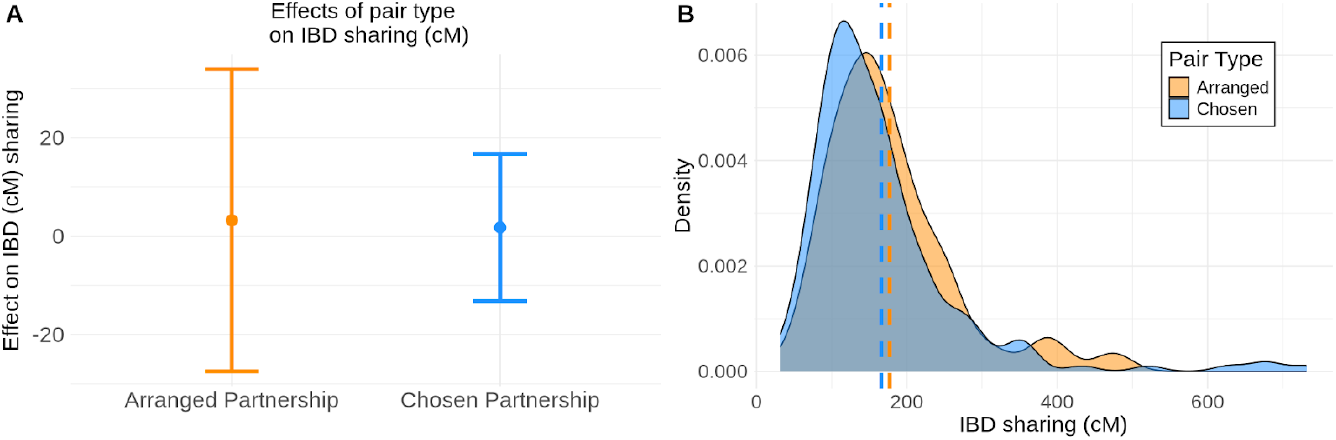
No effect of partnership types on genome-wide IBD sharing. A) We consider a linear model where the intercept is the expected genome-wide IBD sharing based on a background distribution of unrelated female - male pairs from the population. We plot the effect size estimates of arranged partnerships and chosen partnerships on IBD sharing and find that the 95% confidence intervals of the effects contain zero, indicating there is no significant effect of arranged or chosen partnerships on IBD sharing. B) The distributions of the IBD sharing (cM) for arranged and chosen partnerships clearly overlap. Means of each distribution are noted with a dashed line.

### (a) HLA similarity does not differ among partnership types

We next addressed our main question: do couples who have chosen to be together have lower HLA similarity than arranged partnerships? That is, is mate choice driven by minimizing relatedness at the HLA region rather than genome-wide relatedness? We assessed HLA genotype similarity in two ways, first by constructing a HLA similarity score, the number of shared alleles between a pair of individuals across all 8 genes (i.e. minimum score of 0, maximum score of 16) and also by computing the average log odds of homozygous offspring across all 8 genes (see *Methods*). These two metrics are highly correlated (Pearson r = 0.96), but average log odds of homozygous offspring provides a finer measurement (Figure S5). Using linear mixed models for dyadic data that correct for overall IBD sharing and account for the unique influence of each female and male to their similarity scores across all pairings (43) (see eq 3 in *Methods*), we found no effect of arranged or chosen partnership on overall HLA similarity (p = 0.7, 0.61, respectively) (Figure 3A) or on average log odds of HLA homozygosity in potential offspring (p = 0.79, 0.80, respectively) (Figure 3C). In a pairwise contrast, there were no differences between the effects of chosen and arranged partnerships on HLA similarity score (p = 0.56) or on average log odds of HLA homozygosity in potential offspring (p = 0.72). We also assessed whether there was a difference in HLA peptide binding groove protein divergence (exon 2 and 3 for HLA class I genes, exon 2 for HLA class 2 genes) by calculating the grantham distance between the amino acid sequences (33, 34) coded by partners’ alleles. Again, we found no significant effects of arranged or chosen partnerships on mean amino acid sequence divergence between the molecules coded by partners’ HLA genotypes (*p* = 0.74, 0.70, for arranged and chosen partnership effects, respectively, Figure S6) and no significant difference in their effects from a pairwise contrast (*p* = 0.64)

**Figure 3.**
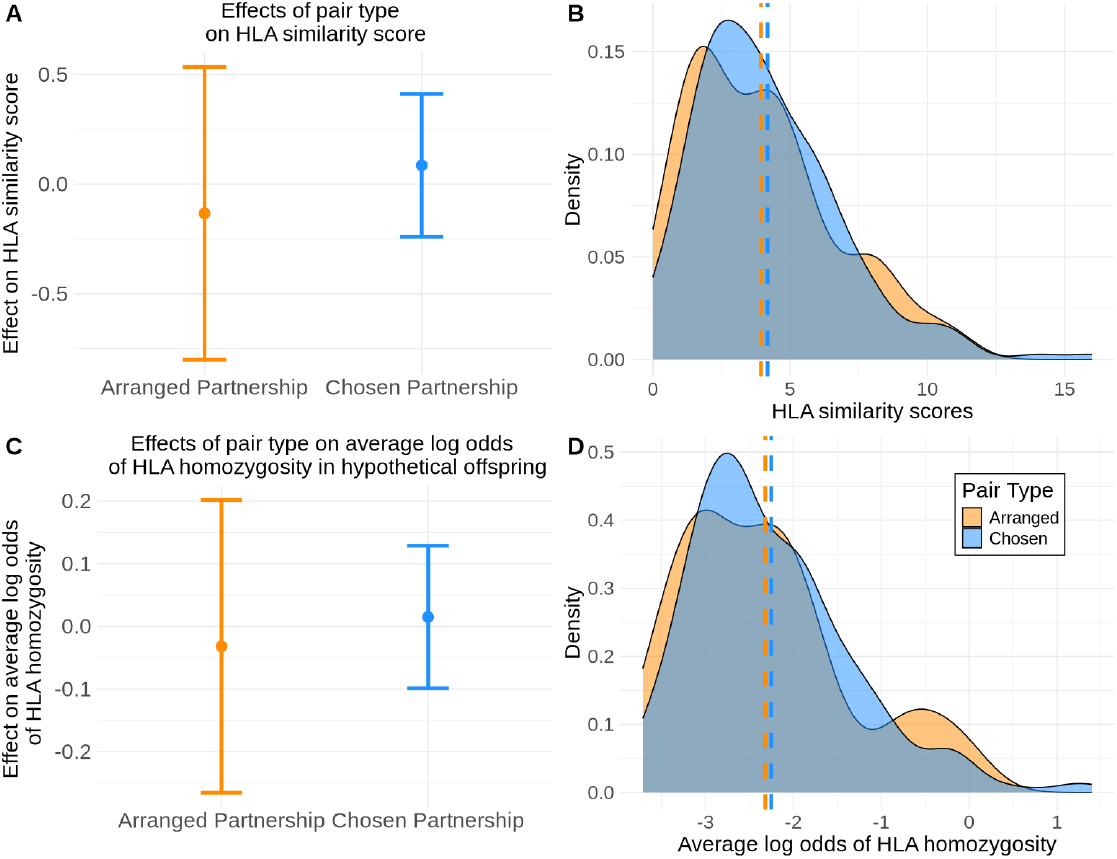
No effect of partnership types on HLA similarity or average log odds of offspring HLA homozygosity. We again consider linear models where the intercept is the expected outcome based on a background distribution of random, unrelated, opposite sex pairs from the population and control for genome-wide IBD sharing. We apply the model to the HLA similarity score (A) and to the average log odds of HLA homozygous offspring (C) across all 8 genes. We plot the effect size estimates of arranged partnerships and chosen partnerships on HLA similarity score (A) and average log odds of offspring homozygosity (C) and find the 95% confidence intervals contain zero, indicating no significant effect of arranged or chosen partnerships on HLA similarity or average log odds of offspring homozygosity. The distributions of HLA similarity scores (C) and average log odds of offspring HLA homozygosity (D) clearly overlap between arranged and chosen partnerships. Means of each distribution are noted with a dashed line.

**Figure 4.**
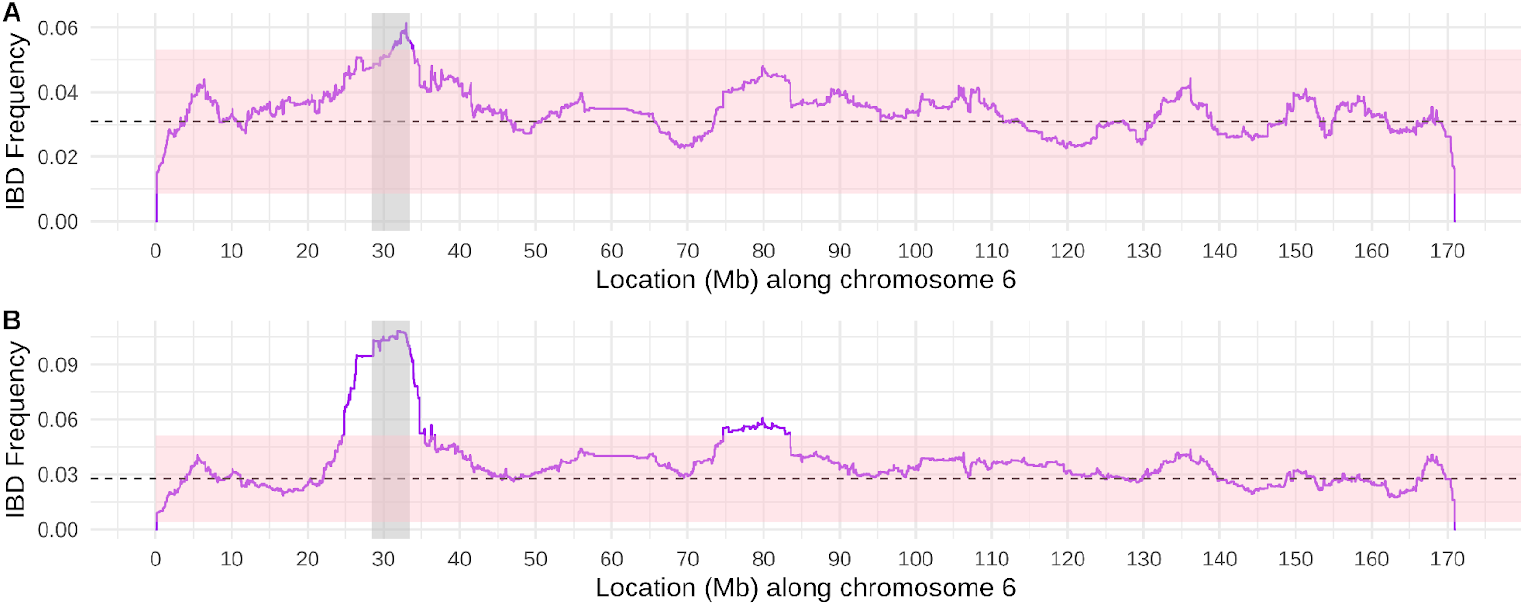
Elevated IBD sharing at the HLA region is consistent with recent, fluctuating positive selection. Panels A and B depict IBD sharing rates across chromosome 6 (rates were averaged within 1000 base pair bins). The dashed line depicts the genome-wide median IBD sharing rate, excluding chromosome 6. Panel A depicts IBD sharing rates for the pairs that make up the background distribution for the partnership analyses (random heterosexual pairings, with relevant age gaps, related at no closer than 3rd degree; n=352 individuals). Panel B depicts the IBD sharing rates for the unrelated individuals included in the allele frequency estimations (related at no closer than 4th degree) with the common haplotypes shown in Figure S9 (n=71 individuals). The HLA region is designated in grey. The pink shaded region depicts +2 and -2 standard deviations from the genome-wide (excluding chromosome 6) median IBD sharing rate, 3.1% and 2.8% for panels A and B, respectively.

### (b) Pathogen binding repertoires of potential offspring do not differ among partnership types

Increased pathogen binding ability of offspring is theorized to be the alleged evolutionary driver of disassortative mating at the MHC region (50, 51). Therefore, even if similarity scores, as assessed by allelic or protein similarity do not differ among pair types, individuals might seek out complementary HLA genotypes in mates that increase the pathogen binding repertoire of potential offspring. Using the same linear mixed model (*Methods* eq 3) as above, we looked for effects of arranged and chosen partnerships on the average computationally-predicted pathogen binding repertoires in hypothetical offspring (i.e., the average number of unique pathogen peptides predicted to be bound by each offspring HLA genotype). We used a curated list of 25 pathogens, taking a subset of those listed in a previous study of evolutionarily significant human pathogens (33) that have plausible transmission in the Himba, and included additional sexually transmitted pathogens known to be prevalent in the Himba (52–54) (personal correspondence with Dr. Ashley Hazel) (see Appendix Table 1). At each locus, the hypothetical offspring genotypes, in expected proportions, were cross-referenced to respective binding affinities to our list of antigenic peptides producing a mean number of unique pathogen peptides that strongly bind per HLA genotype. Many of the HLA class II DPA/B or DQA/B gene products can form both cis and trans heterodimers, comprising DP and DQ epitopes, respectively (39, 40). Thus, an individual heterozygous for both DPA1 and DPB1 may form 4 unique DPα1/DPβ1 molecules. We assessed the mean number of bound antigenic peptides for all possible offspring DP and DQ epitope combinations, again based on expected genotype proportions.

Across all 8 HLA genes tested, there was no effect of arranged or chosen partnerships on the mean number of pathogen peptides bound by offspring allotypes that differed from the background distribution of random partners (Figure S7, Table S4). Likewise, we found no differences in the effect estimates of arranged and chosen partnerships in pairwise contrasts (Table S5). HLA class I molecules generally bind peptides derived from intracellular pathogens (e.g. viruses) whereas HLA class II molecules generally bind those derived from extracellular pathogens (e.g. bacteria and parasites) (55). Thus, we also performed this analysis including only intracellular pathogens for HLA-A, B, and C genotypes and only extracellular pathogens for HLA-DP, DQ, and DR genotypes. We still found no significant effects of arranged or chosen partnerships in mean pathogen peptides bound and no significant differences of effects in pairwise contrasts (Figure S8, Tables S6-7).

### (c) Extensive haplotype sharing could indicate recent episodes of fluctuating positive selection

We find no evidence that disassortative mating is driving maintenance of polymorphism at the HLA. Other forms of selection, besides sexually mediated balancing selection, could therefore play a larger role in maintaining polymorphism at this locus. We assessed haplotype sharing by visualizing the SNP-based haplotypes (from genotyping array) across the entire 5Mb region surrounding the 8 genotyped HLA genes. The SNP-based haplotypes along this much larger 5Mb region were grouped by their HLA allele haplotypes (typed via short-read sequencing) (Figure S9). Although the 7 HLA genes used to create allele-based haplotypes comprise only ∼1% of the base pairs in this 5Mb region, we see virtually identical SNP haplotypes across unrelated individuals (maximum relatedness of 4^th^ degree) who share frequent HLA allele-based haplotypes (n=92 haplotypes with >4 occurrences in the 102 unrelated individuals) (Figure S9-10). This finding indicates that the entire 5Mb HLA region was inherited without recombination in these unrelated individuals, consistent with recent positive selection on these haplotypes (56, 57).

Unlike our previous study of sub-Saharan African populations, where in 6 of the 7 populations the majority of haplotypes were observed only once (45), 70% of haplotypes were observed at least twice in our set of unrelated individuals, with the most frequent haplotype (*A*30:02 - C*03:04 - B*15:10 - DRB1*03:01 - DQA1*05:01 - DQB1*02:01 - DPB1*01:01*) observed 23 times in the unrelated set. Nineteen of the 29 haplotypes occurring more than once in the unrelated set of individuals were observed in at least one of the 7 sub-Saharan African populations from our previous study (Figure S11). We assessed whether there was a relationship between haplotype frequency and number of unique peptides computationally predicted to be bound by each haplotype (58), but found no significant relationship (*p* = 0.91, Figure S12) and likewise found no significant relationship with number of peptides bound when we analyzed each pathogen independently (Figure S13).

Consistent with the SNP-based haplotype visualization, we found increased IBD sharing at the HLA region relative to genome-wide levels. This pattern implies recent (within 200 generations) positive selection as opposed to overdominance (i.e., heterozygote advantage) (56, 57). The mean rate of IBD sharing across the HLA region between random pairs of unrelated individuals from the background distribution used in our pair analyses (n=352 individuals, n=18,487 pairs, related at a maximum of 3rd degree) is 5.3%, 1.97 standard deviations above the genome-wide median of 3.1%. The rate of IBD sharing across the HLA region in the subset of unrelated individuals from our allele frequency analyses (related at a maximum of 4th degree) who have common HLA-based haplotypes (n=71 individuals, haplotypes shown in figure S9) is even higher at 10.4%, 6.4 standard deviations above the genome-wide median of 2.8%.

## 4. Discussion

This study contributes an important new data point to the growing literature on HLA and mate choice. Unlike studies based on cosmopolitan datasets—where apparent HLA similarity in couples may arise spuriously from preferences for ethnically, and therefore genetically, similar partners—our study examines a small-scale population in which such confounding is minimized. Small-scale societies like the Himba more closely approximate evolutionarily relevant social contexts, characterized by high endogamy, which may make any preferences for HLA dissimilarity more relevant that in societies where any unrelated individual is sufficiently dissimilar at their HLA. To date, only one other relatively small-scale population (European Americans from the Mormon community (7, 15)) has been examined with comparable genome-wide relatedness measures.

Cultural norms like arranged marriage and cross-cousin marriage preferences are predicted to promote similarity across the HLA, while tolerance of extramarital partnerships and “love match” marriages could enhance the possibility of seeking out an HLA-dissimilar partner. Therefore, the unique socioecology of Himba marriage patterns sets up a strong test of the dissimilarity hypothesis–contrasting HLA similarity in arranged and chosen partnerships.

Surprisingly, we find that chosen partnerships are neither less similar at their HLA than random pairs constructed from the population background distribution, nor do they differ in their HLA similarity from arranged partnerships. We also find no evidence that chosen partners are selected to directly maximize an offspring pathogen binding repertoire. Unlike studies of Hutterites (6) and Mormons (7) that do show evidence for greater HLA dissimilarity in married pairs, background genetic diversity in an African population like the Himba might be high enough that any unrelated individual is sufficiently dissimilar at the HLA and there may be no fitness benefit to maximizing dissimilarity. It has been suggested that offspring may not benefit from maximal diversity at the HLA; T cells that bind self peptide-MHC complexes too strongly undergo pruning in the thymus. Possessing many different MHC alleles could deplete the T cell repertoire required for appropriate immune response (59, 60). This process of negative selection for extremely autoreactive T-cells has been hypothesized to inhibit the expansion of the MHC genomic region to include additional copy numbers (61). Supporting this optimality hypothesis, decreased parasite load in intermediate versus maximally diverse MHC individuals has been observed in voles (60) and stickleback fish (62) and our data suggest that this may also be relevant for humans.

We find extensive IBD sharing at the HLA region consistent with recent, fluctuating positive selection which may contribute to polymorphism maintenance at this region, rather than heterozygote advantage (56, 57). Despite association with many diseases (63) and consistently showing up as a target of natural selection (58, 61), the selective forces shaping HLA diversity are not well understood (61). We suggest that innate drive for disassortative mating based on HLA genotype is unlikely to be a major evolutionary force in maintaining diversity in evolutionarily relevant social contexts. Future work should focus on assessing the contribution of other selective regimes (e.g., negative frequency dependent, fluctuating selection, heterozygote advantage, divergent allele advantage, etc.) to the maintenance of polymorphism at the HLA region.

Assortative mating for sociocultural traits may be a more important driver of mate choice for groups like the Himba rather than maximizing HLA diversity. Our previous work has shown that resource scarcity affects women’s partner preferences (64, 65), and that relationship success is correlated with similarity in mate value (66). Concurrent partnerships, rather than increasing the chance of finding a partner with a dissimilar HLA genotype, may be more important for garnering resource security for women, and distributing reproductive opportunities for men who are otherwise limited by polygyny and a later age at marriage (17, 67).

## Ethics

Ethical approval for this study was granted by the University of California, Los Angeles (IRB-10-000238), the State University of New York, Stony Brook (IRB-636415-12), and was approved by the Namibian Ministry of Home Affairs and the University of Namibia Office of Academic Affairs and Research. Chief Basekama Ngombe provided permission to work in the community and local approval of the study. Community leaders were actively involved in discussions regarding what genetic data could be used for, who would have access to it, and whether there was a for-profit element involved (there was not). Individual informed consent, and for minors’ parental assent, was obtained orally from all participants, given low rates of literacy in the community. Care was taken to protect participants’ privacy, for example via a double-blind procedure for DNA collection (68). These data were collected as part of the longitudinal Kunene Rural Health and Demography Project, which has been working in the community since 2010.

## Acknowledgements

We thank the Himba communities in which we have worked; without their support, this study would not have been possible. We thank Dr. Mark Grote and Cole Williams for assistance with statistics and data analysis. We thank Dr. Ashley Hazel for guidance in selecting ecologically relevant pathogens. BMH and GLM were supported by the National Institutes of Health (NIH) grant R35GM133531. BS and SP were supported by the National Science Foundation grant BCS-1534682. BMH, GLM, TDJF, NFP, and PJN were supported by NIH award R01AI151549. The authors acknowledge the High Performance Computing Core Facility at the University of California, Davis for providing computational resources that have contributed to the research results reported in this paper. The content is solely the responsibility of the authors and does not necessarily represent the official views of the National Institutes of Health or the National Science Foundation. The funders had no role in the decision to publish or prepare the manuscript.

## Author Contributions

BS, PN, and BMH conceived of the study. GLM conducted analyses with help from KH and CB. GLM and BS wrote the manuscript with input from PN, SP, and BMH. KK, GH, NFP, TDJF, and NP generated HLA sequence data under the supervision of PN. Sample and ethnographic data collection were conducted by BS and SP. BMH and BS supervised the research.

## Use of AI-assistance

AI was not used in the preparation of this article.

## Data and code availability

Genotype and HLA allele call data for the individuals included in this study have been deposited at dbGap as phs001995.v3.p1. All original code and data used to conduct analyses are deposited in the following github repository https://github.com/gillianmeeks/Himba_HLA_mating and on dryad https://doi.org/10.5061/dryad.02v6wwqg2. Partnership type assignments are not available due to the sensitive nature of the data.

## Supplemental information

SI. Supplementary Figures S1–S13 and Supplementary Tables S1–S7

Appendix Table 1. Spreadsheet listing pathogen peptide NCBI accessions, related to methods.

