## Supplementary Figures and Tables for "No evidence for disassortative mating based on HLA in a small-scale, endogamous population"

[9] Lead contact

**This PDF file includes:**
Supplementary Figures S1–S13
Supplementary Tables S1-S7

**Other supporting materials for this manuscript include the following:**
Appendix Table 1

**Figure S1:** Asymmetric Linkage Disequilibrium (LD) statistics based on our phased HLA-allele based haplotypes. Asymmetric LD takes into account the differences in the number of alleles between the two loci. The values in each square are the LD scores of the row's gene name conditional on the number of alleles present at the column gene.

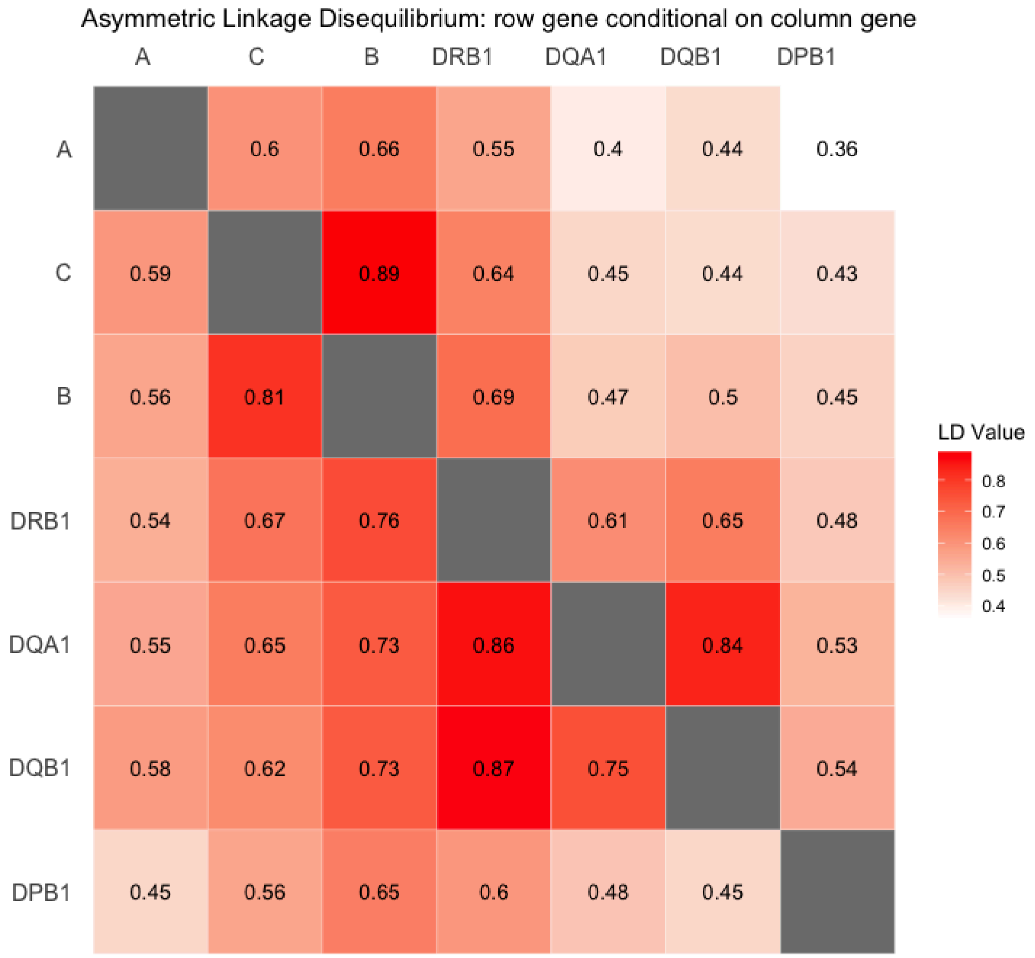

**Figure S2:** Depiction of the line sweep algorithm used to calculate IBD sharing rates genome-wide. A) The range is marked for each IBD segment in a dataset. B) The markings are tallied within a newly generated chromosome length array. The ‘start’ of an IBD segment is marked +1, and the ‘end’, the base pair directly following the last base pair, is marked -1. C) A ‘line’ sweeps across the chromosome-length array carrying over tallies from the previous base pair to generate IBD coverage scores for each base pair.

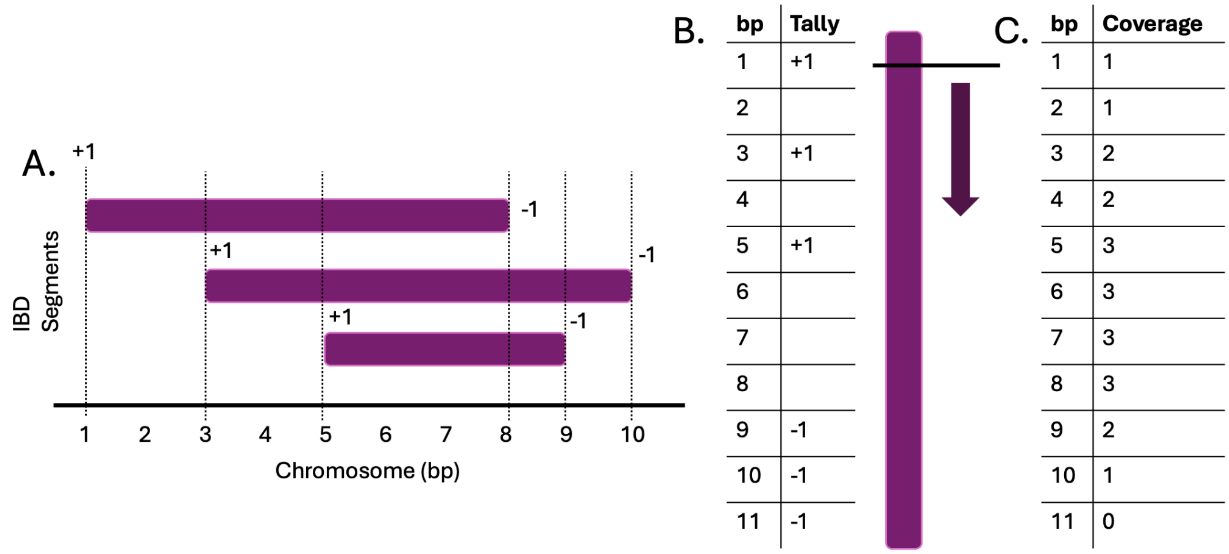

**Figure S3:** Allelic diversity at the 8 HLA short read sequenced genes (A, B, C, DPA1, DPB1, DQA1, DQB1, DRB1) across the full sample (n=366 Individuals), which includes relatives. We assessed whether any alleles that were present in the Himba were rare or absent in the rare allele database (1). Two alleles were determined to be globally rare, DPA1*01:05, present in only 5 copies in the database (4 copies in the Spanish Basque, and 1 copy in a United Arab Emirates population), and DPB1*61:01N, present in only 2 copies (1 copy in a South African and 1 copy in the Cameroonian Saa).

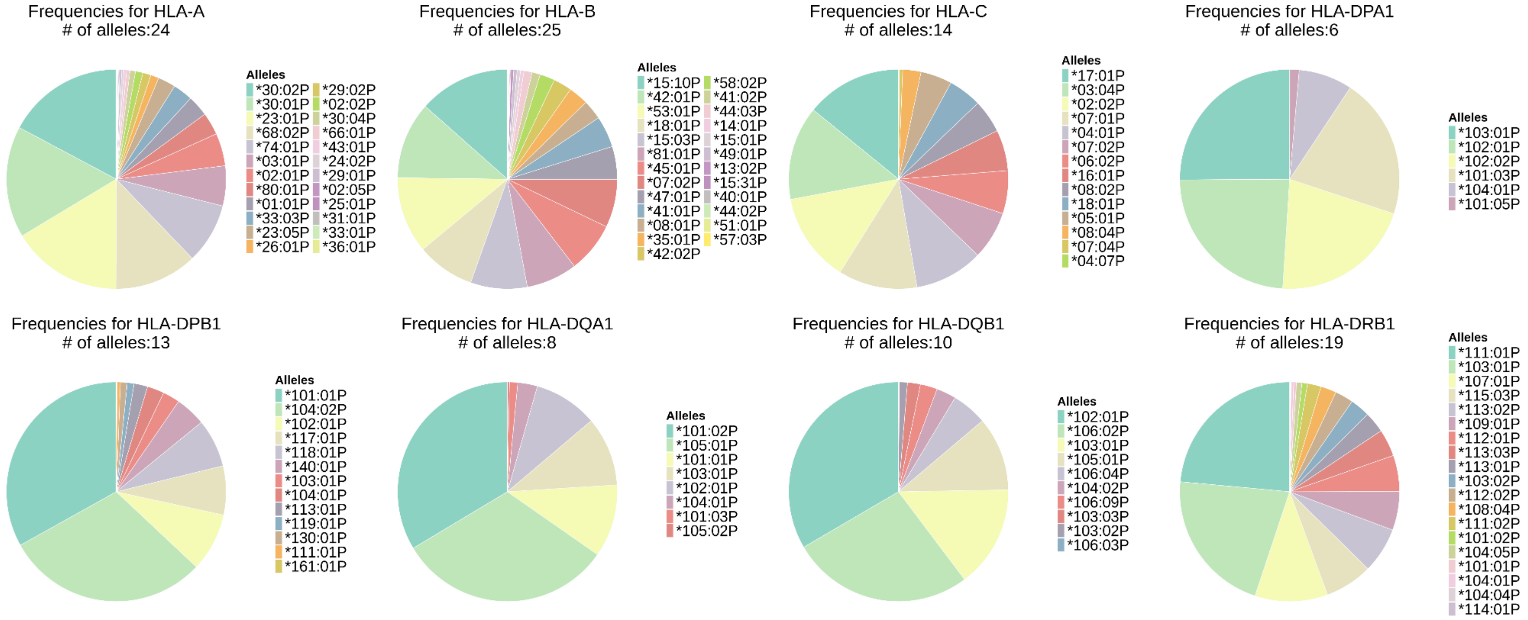

**Figure S4:** Allelic diversity at the 8 HLA short read sequenced genes (A, B, C, DPA1, DPB1, DQA1, DQB1, DRB1) across the maximum set of unrelated individuals (n=102), closest relation allowed is 4th degree.

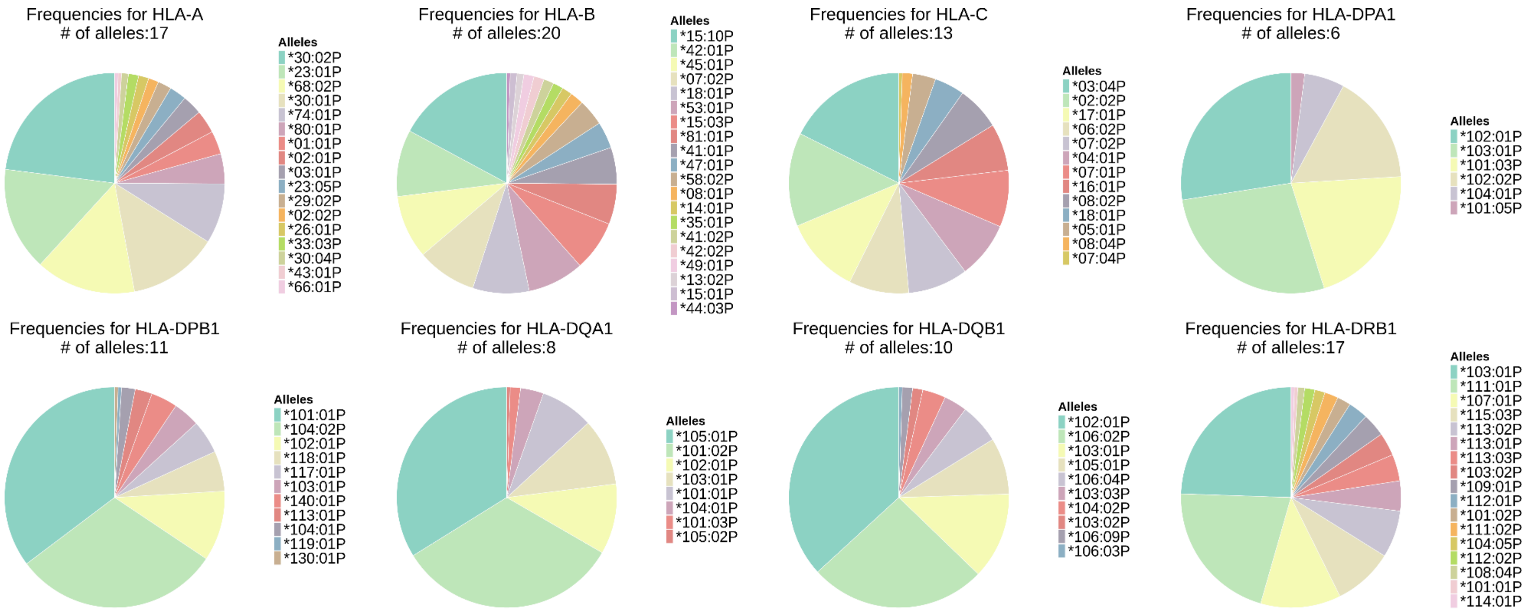

**Figure S5:** Relationship between HLA similarity score and average log odds of HLA homozygosity in offspring in the full dataset used to construct models, including the background distribution. HLA similarity score and average log odds of HLA homozygosity are highly correlated (Pearson r = 0.96).

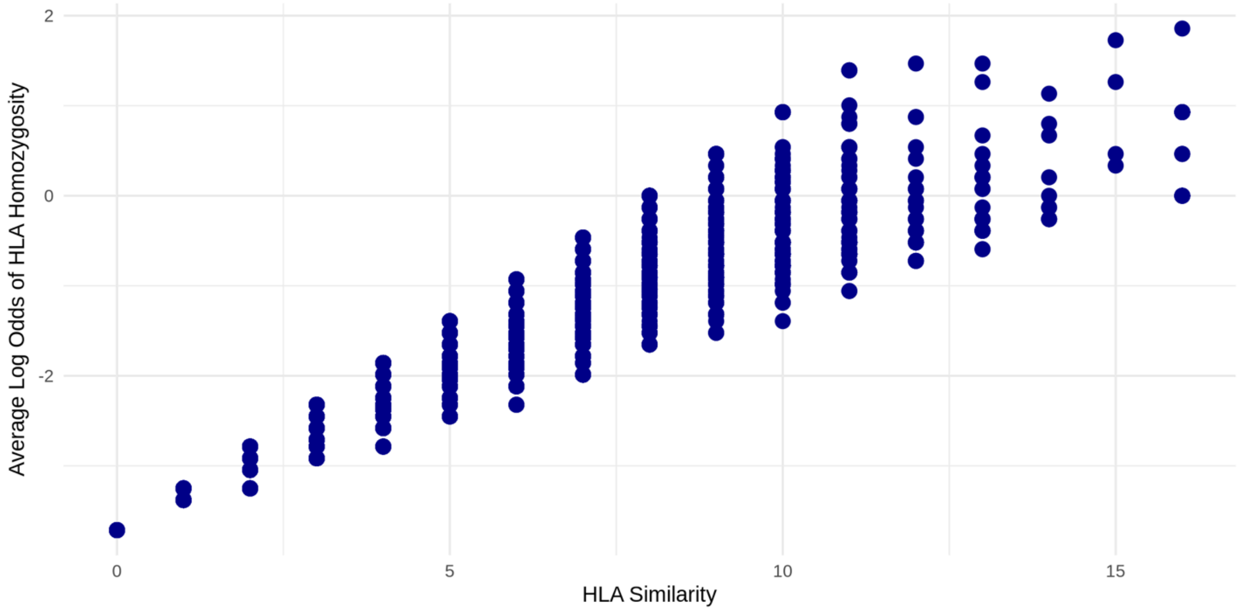

**Figure S6:** No effect of arranged or chosen partnerships on mean peptide binding groove sequence divergence across all 8 HLA genes. Left column shows effect size estimates of the partnership types and the right column shows the distributions of the mean sequence divergence scores across all 8 genes between arranged and chosen partnerships.

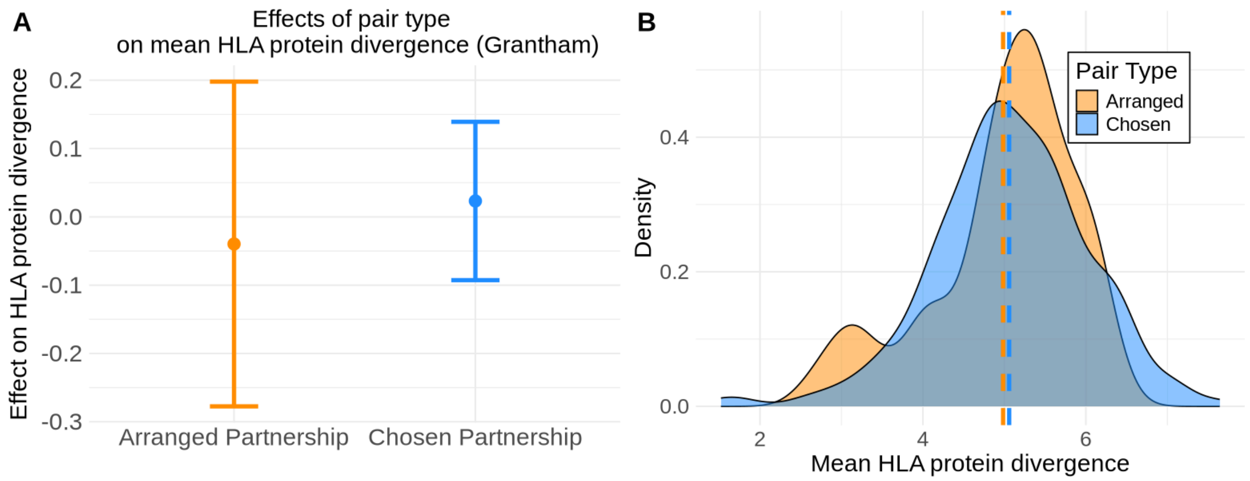

**Figure S7:** No effect of arranged or chosen partnerships on pathogen binding scores (mean number of pathogen peptides bound across potential offspring genotypes) at any of the HLA genes. Left column shows effect size estimates of the partnership types and the right column shows the distributions of the number of bound pathogen peptides based on potential offspring genotypes between arranged and chosen partnerships. Gene products are ordered HLA-A, HLA-B, HLA-C, HLA-DP, HLA-DQ, and HLA-DRB1 from the top row to bottom row.

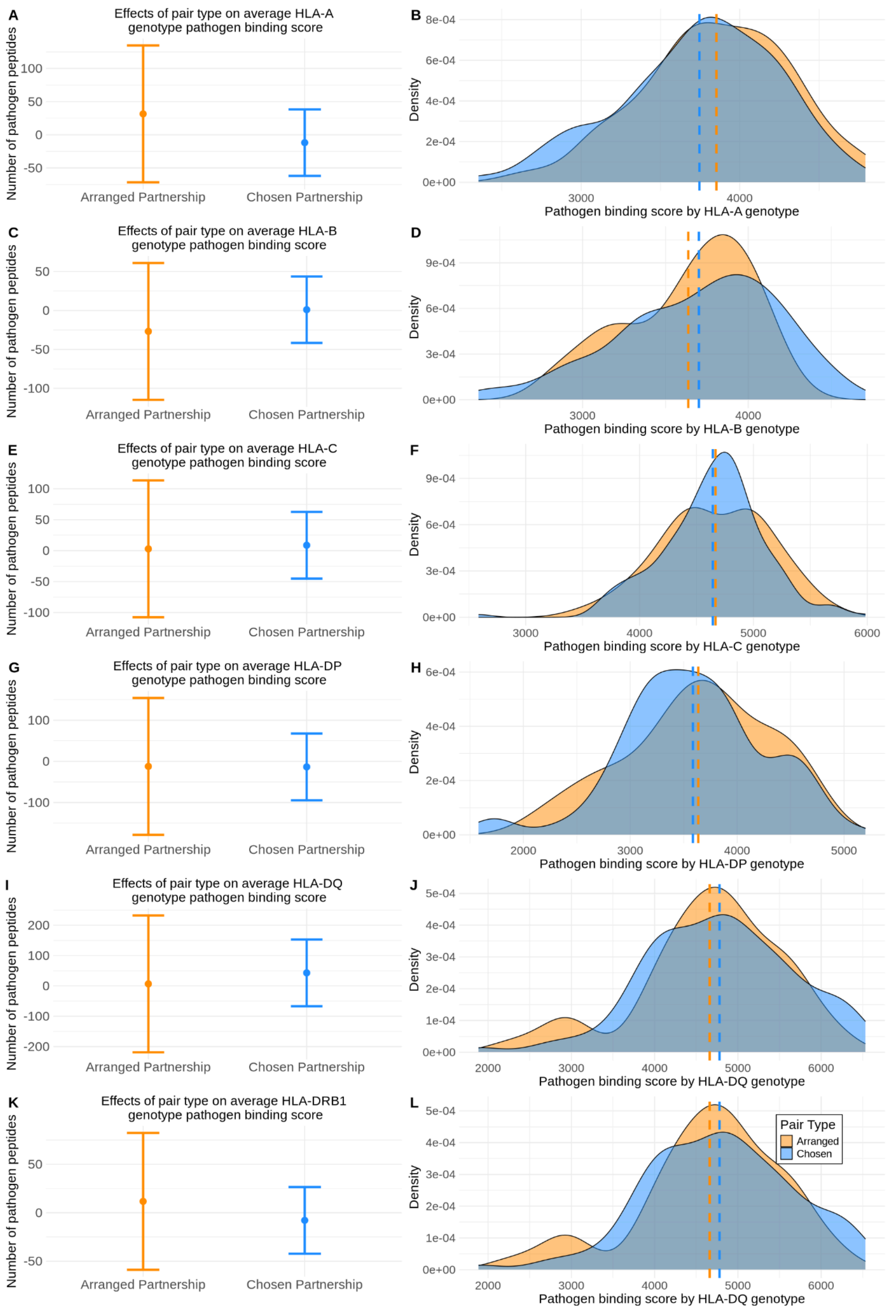

**Figure S8:** No effect of arranged or chosen partnerships on pathogen binding scores (mean number of pathogen peptides bound across potential offspring genotypes) at any of the HLA genes for class relevant pathogens. Peptide lists were restricted to functionally relevant pathogens (intracellular pathogens for HLA class 1 genes and extracellular pathogens for HLA class 2 genes). Left column shows effect size estimates of the partnership types and the right column shows the distributions of the number of bound pathogen peptides from potential offspring between arranged and chosen partnerships. Gene products are ordered HLA-A, HLA-B, HLA-C, HLA-DP, HLA-DQ, and HLA-DRB1 from the top row to bottom row.

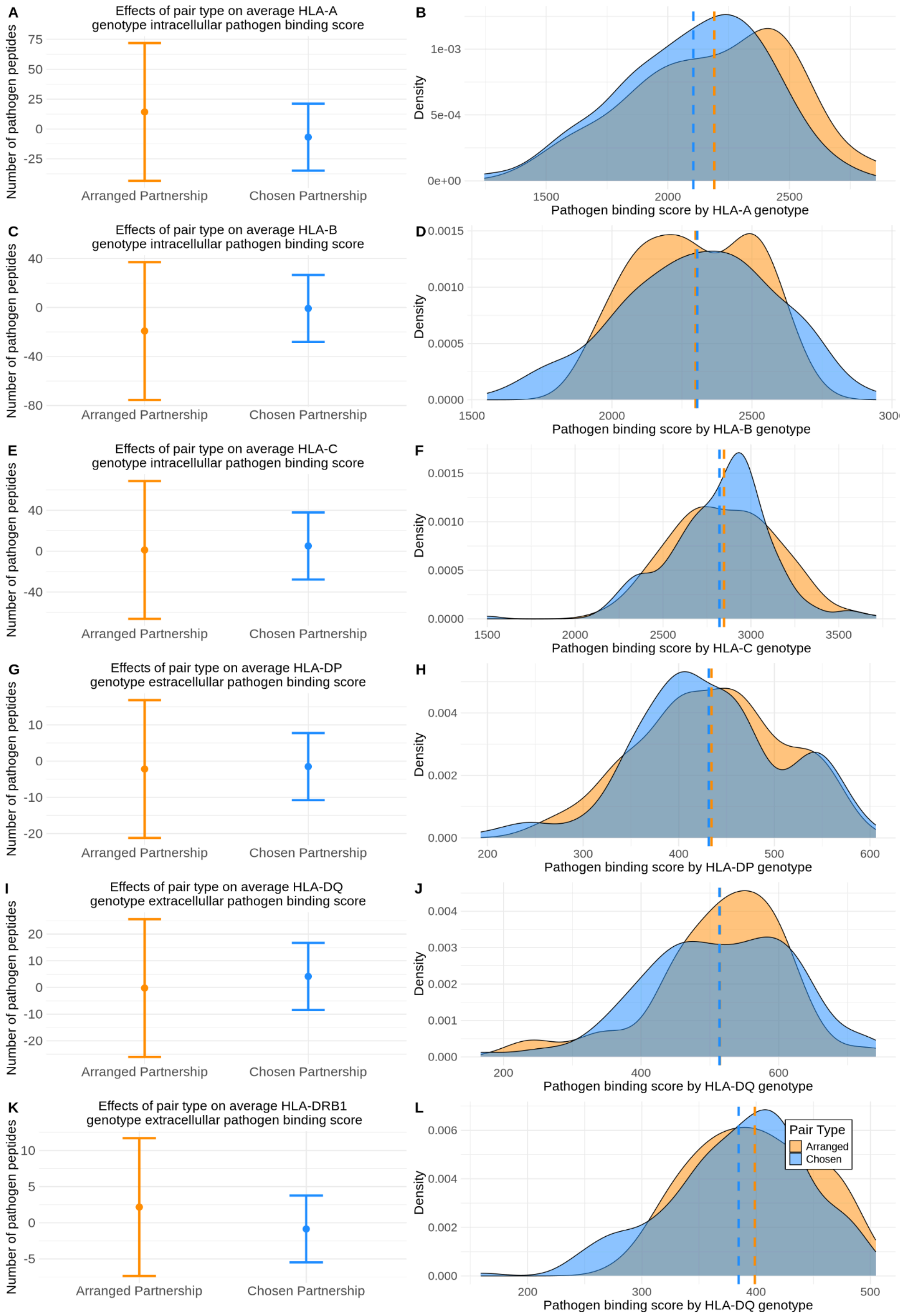

**Figure S9:** *Extensive haplotype sharing across the 5 Mb HLA region among unrelated individuals*. SNP-based haplotypes (rows) are grouped according to HLA-allele based haplotypes (for those haplotypes with > 4 occurrences in the 102 unrelated individuals). Columns indicate genomic positions with white/black designating the ref vs. alt allele at each position. The haplotypes are grouped by their allele calls at HLA-A - C - B - DRB1 - DQA1 - DQB1 - DPB1. We were unable to resolve the haplotypes for DPA1 alleles calls. Haplotypes were visualized with haplostrips (2). The approximate, relative locations of the 7 HLA genes are shown on the x-axis along with the approximate starting and ending positions of the upstream Olfactory Receptor (OR) gene cluster.

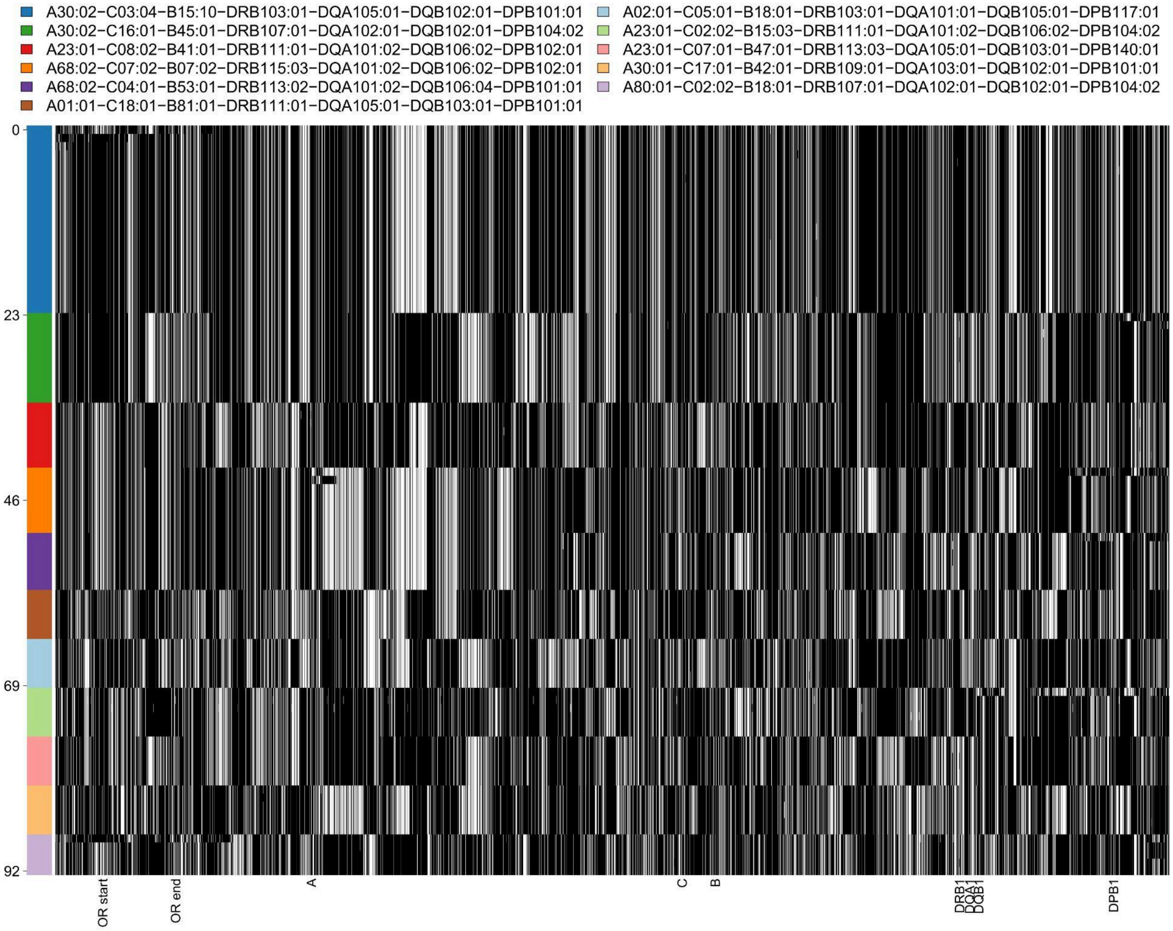

**Figure S10:** Number of SNP differences in a 5Mb window (from genotype array data) across the common (>4 occurrences) HLA allele-based haplotypes relative to the most frequent haplotype (index 0, bottom row) are shown. This figure quantitatively depicts the number of SNP differences across the haplotypes shown in Figure 4 of the main text. Allele-designated haplotypes are largely identical by state as evident by the clustering of haplotypes with the same number of differences to the reference haplotype.

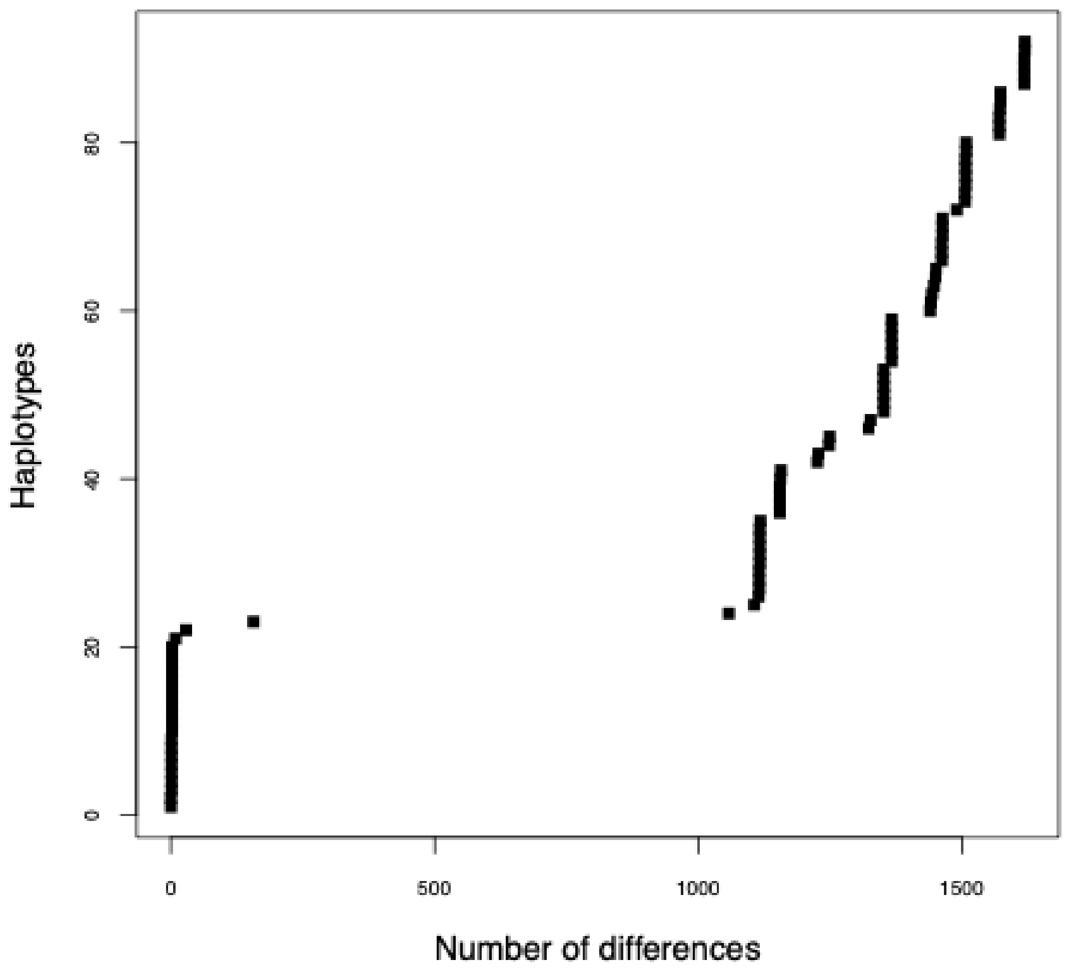

**Figure S11:** Common (>1 occurrence in the unrelated set of 102 Himba individuals) class I haplotype frequencies as compared to frequencies in 7 other sub-Saharan African populations (3). Himba frequencies are in red. 10 of the 29 common class I haplotypes are found only in the Himba and not in the other 7 sub-Saharan African populations.

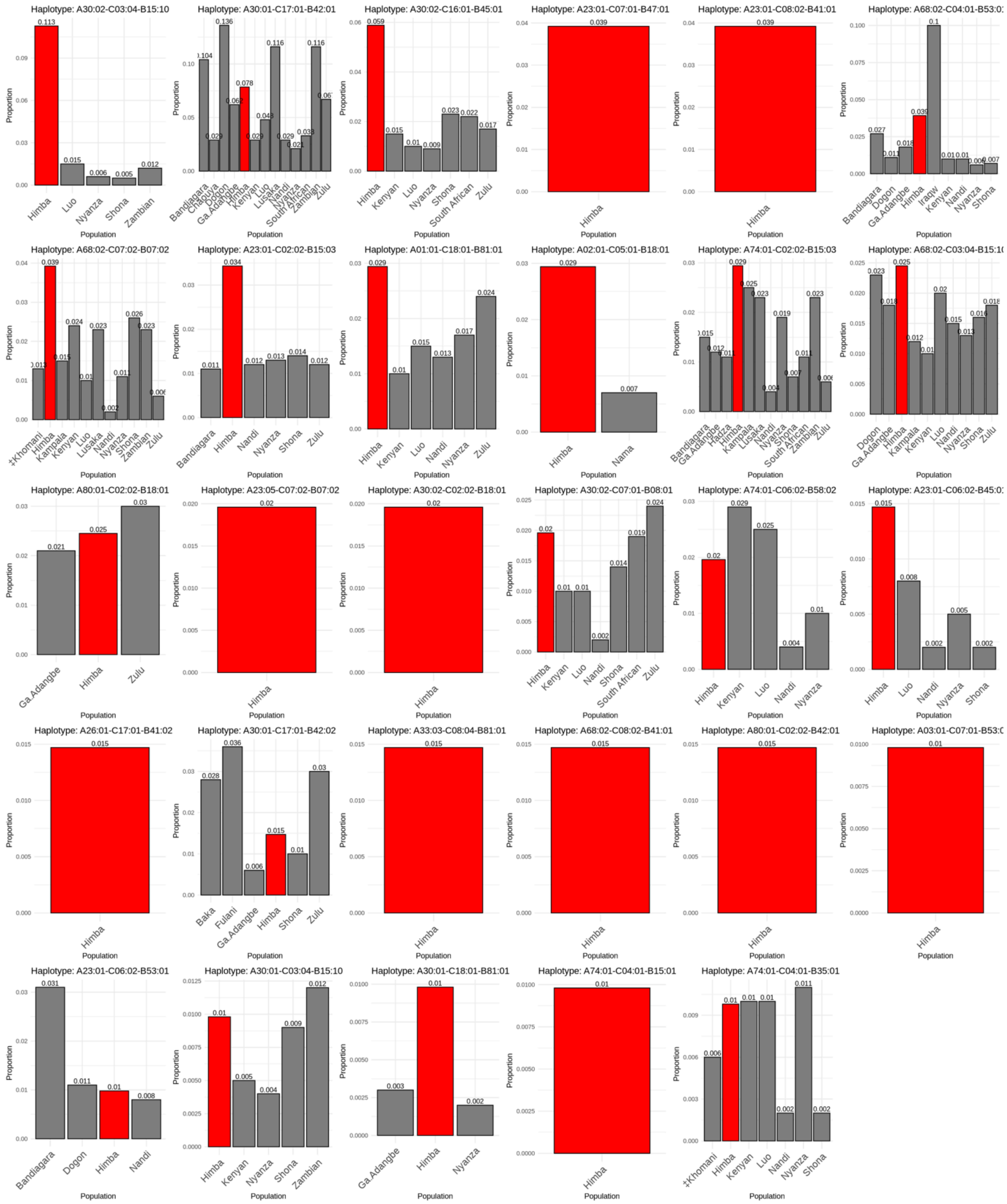

**Figure S12:** No relationship between the frequency of common (>1 occurrence in the unrelated set, n=102 Himba individuals) haplotypes with the number of unique pathogen peptides they are predicted to bind across all 25 pathogens.

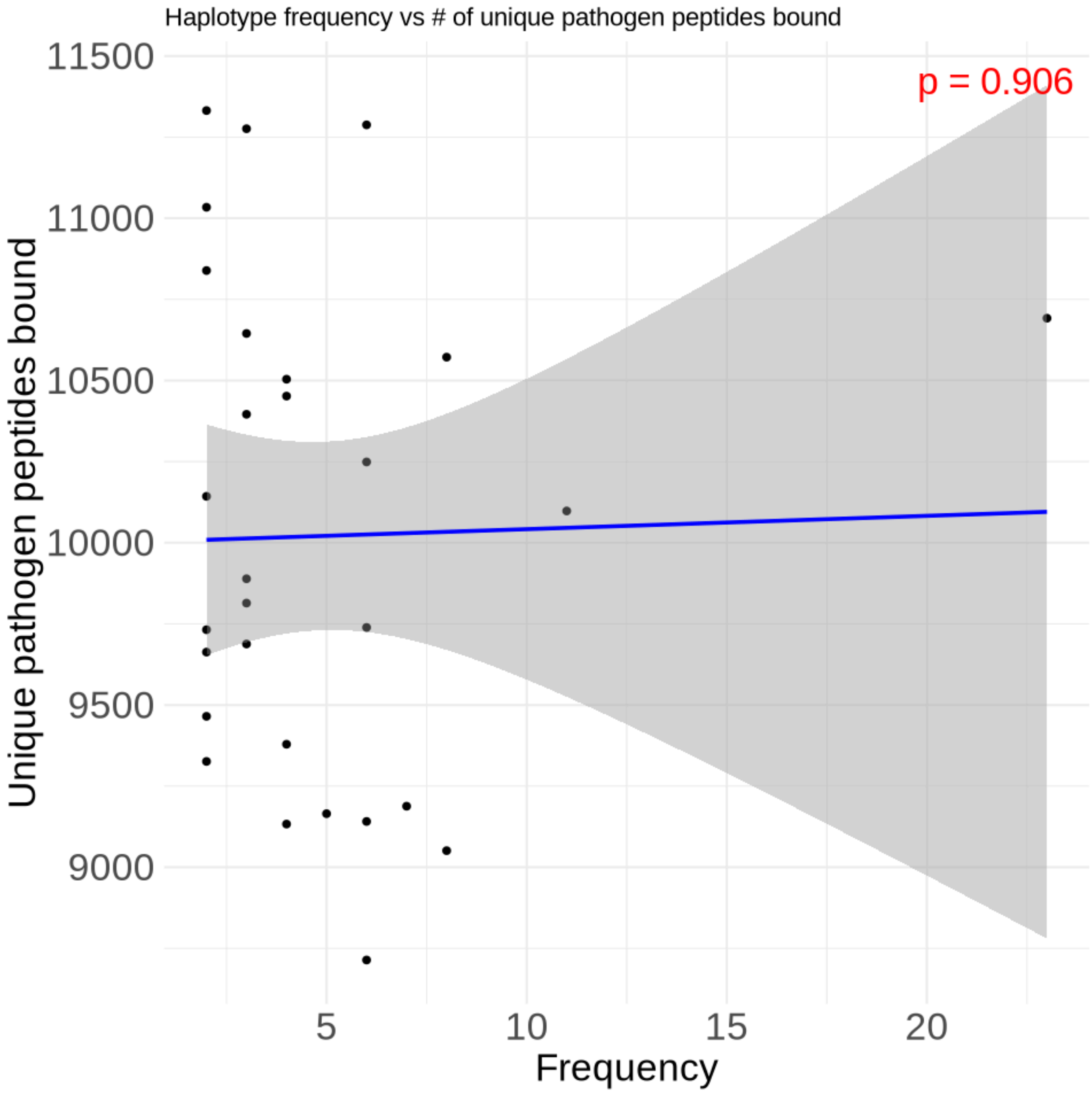

**Figure S13:** No relationship between the frequency of common (>1 occurrence in the unrelated set, n=102 Himba individuals) haplotypes with the number of unique pathogen peptides predicted to be bound, for any pathogen.

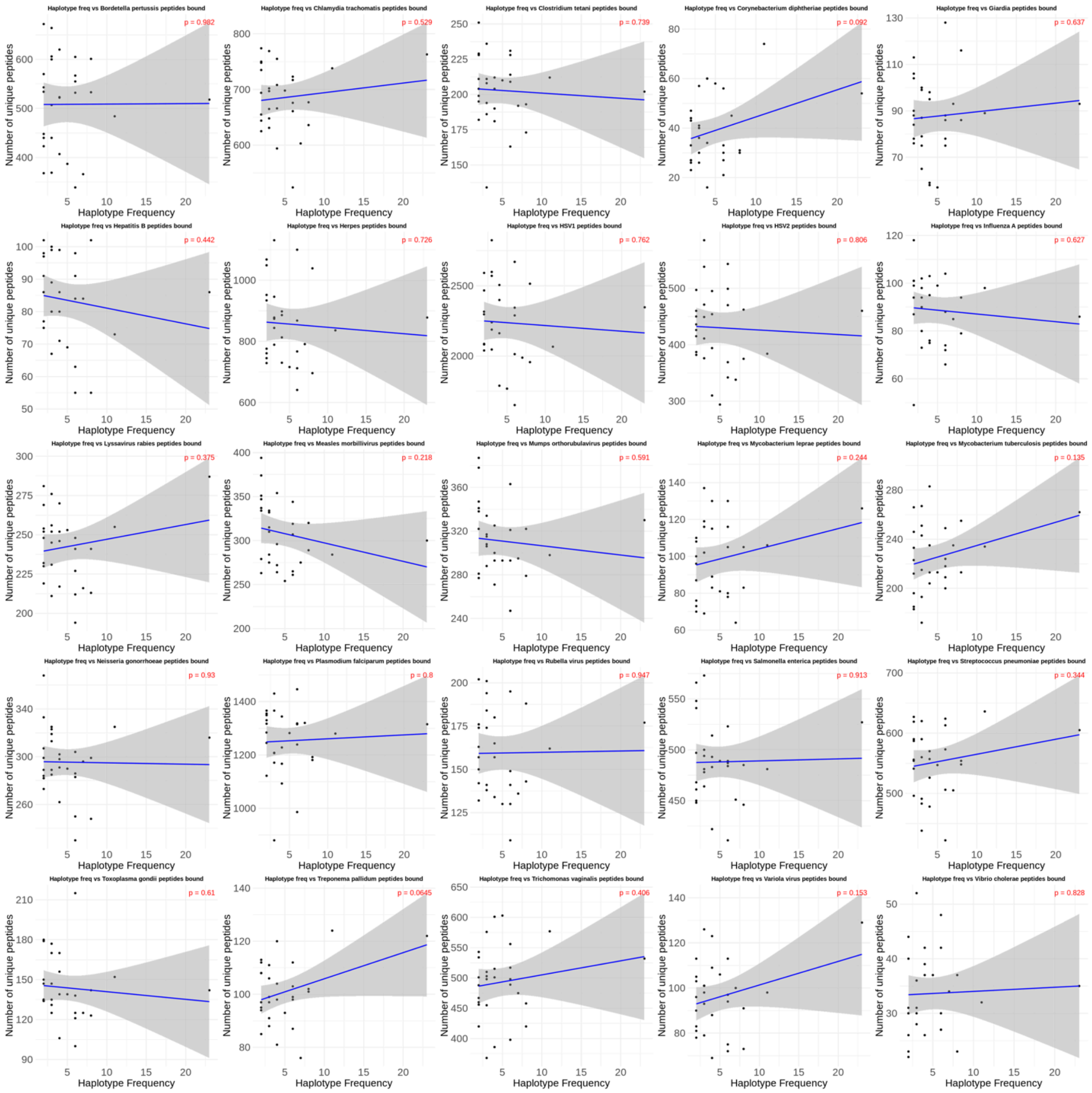

**Table S1:** IBD calling accuracy across 4 software program runs

| | **Germline 1.5.3** | **Germline 2** | **PhasedIBD (phase correction = TRUE)** | **PhasedIBD (phase correction = FALSE)** |
|---|---|---|---|---|
| Average unrelated pairs IBD (cM) | 131 | 134 | 161 | 154 |
| Average parent child IBD (cM) | 2551 | 2526 | 2978 | 2580 |
| Total map length (cM) | 2472 | 2472 | 2472 | 2472 |
| Normalized parent child sharing (percent of map length) | .979 | .968 | 1.14 | .981 |

Normalized parent-child IBD sharing was calculated by subtracting the average unrelated pairs (> 3rd degree relation) IBD sharing, an estimate of background IBD sharing in the population, from the average parent-child IBD sharing and normalizing by the map length. Assuming the average unrelated IBD sharing accurately represents IBD2 shared by parent child pairs, then the remaining IBD sharing should span the entire map length, i.e., the whole genomic copy inherited from parent to child and the normalized value should be close to 1. The parameters used for germline 1.5.3 were -haploid -w_extend and for GERMLINE2 were -h -g 2. Default parameters were used for phasedIBD runs. A threshold of 3cM was used for all runs.

**Table S2**: Possible states of odds of homozygous offspring at one locus

| **Number of shared alleles by pair** | **Partner 1 genotype** | **Partner 2 genotype** | **Odds of homozygous offspring** $(\frac{\# \text{ events homozygous}}{\# \text{ events heterozygous}})$ |
|---|---|---|---|
| 2 | Homozygous (e.g., A1/A1) | Homozygous (e.g., A1/A1) | $\frac{4}{0}$ |
| 2 | Heterozygous (e.g., A1/A3) | Heterozygous (e.g., A1/A3) | $\frac{2}{2}$ |
| 1 | Homozygous (e.g., A1/A1) | Heterozygous (e.g., A1/A3) | $\frac{2}{2}$ |
| 1 | Heterozygous (e.g., A1/A3) | Heterozygous (e.g., A1/A2) | $\frac{1}{3}$ |
| 0 | Homozygous or Heterozygous (e.g., A1/A1 or A1/A2) | Homozygous or Heterozygous (e.g., A3/A3 or A3/A4) | $\frac{0}{4}$ |

For a given number of shared alleles (1 or 2) there are multiple possible odds of homozygous offspring depending on if the individuals in the pair are heterozygous or homozygous at the locus.

**Table S3:** No deviations from Hardy Weinberg Equilibrium at the 8 typed HLA genes.

| **Gene** | **Expected Heterozygosity** | **Observed Heterozygosity** | **P-value (Chi-Squared Test)** |
|---|---|---|---|
| HLA-A | 0.870 | 0.863 | 0.998 |
| HLA-B | 0.914 | 0.951 | 0.400 |
| HLA-C | 0.896 | 0.922 | 0.792 |
| HLA-DPA1 | 0.775 | 0.775 | 1.000 |
| HLA-DPB1 | 0.762 | 0.745 | 0.990 |
| HLA-DQA1 | 0.750 | 0.814 | 0.993 |
| HLA-DQB1 | 0.768 | 0.843 | 0.872 |
| HLA-DRB1 | 0.861 | 0.892 | 0.999 |

We ran the ASTA (Asymptotic Statistical Test with Ambiguity) program, from the package HWETests, designed to test for deviations from HWE at polymorphic loci, on the maximized set of unrelated individuals (n=102) (4). Here we show the differences between the expected heterozygosity and the observed heterozygosity along with the Chi-squared test p-value for each HLA gene.

**Table S4**: P-values for the partnership type effects on mean number of pathogen peptides bound by potential offspring genotypes.

| **Outcome Variable** | **Arranged effect p-value** | **Chosen effect p-value** |
|---|---|---|
| A genotype binding | .55 | .65 |
| B genotype binding | .55 | .96 |
| C genotype binding | .96 | .75 |
| DP genotype binding | .89 | .75 |
| DQ genotype binding | .95 | .44 |
| DRB1 genotype binding | .75 | .66 |

P-values shown here correspond to the effect size estimates shown in the left panel of supplementary figure 5.

**Table S5:** P-values for the pairwise contrasts of partnership type effects on mean number of pathogen peptides bound by potential offspring genotypes.

| **Outcome Variable** | **Arranged vs chosen effect p-value** |
|---|---|
| A genotype binding | .46 |
| B genotype binding | .57 |
| C genotype binding | .93 |
| DP genotype binding | .99 |
| DQ genotype binding | .78 |
| DRB1 genotype binding | .62 |

**Table S6:** P-values for the partnership type effects on mean number of biologically relevant pathogen peptides bound by potential offspring genotypes.

| Outcome Variable | Arranged effect p-value | Chosen effect p-value |
|---|---|---|
| A genotype binding | .63 | .63 |
| B genotype binding | .51 | .96 |
| C genotype binding | .978 | .76 |
| DP genotype binding | .82 | .75 |
| DQ genotype binding | .99 | .52 |
| DRB1 genotype binding | .66 | .72 |

Only intracellular pathogens were considered for class I genes (A, B, C) and only extracellular pathogens were considered for class II genes (DP, DQ, DRB1). P-values shown here correspond to the effect size estimates shown in the left panel of supplementary figure 6.

**Table S7:** P-values for the pairwise contrasts of partnership type effects on mean number of biologically relevant pathogen peptides bound by potential offspring genotypes.

| Outcome Variable | Arranged vs chosen effect p-value |
|---|---|
| A genotype binding | .52 |
| B genotype binding | .56 |
| C genotype binding | .92 |
| DP genotype binding | .95 |
| DQ genotype binding | .77 |
| DRB1 genotype binding | .57 |

Only intracellular pathogens were considered for class I genes (A, B, C) and only extracellular pathogens were considered for class II genes (DP, DQ, DRB1).

**Supplementary Figures/Tables References**


1. Gonzalez-Galarza FF, McCabe A, Santos EJMD, Jones J, Takeshita L, Ortega-Rivera ND, et al. Allele frequency net database (AFND) 2020 update: gold-standard data classification, open access genotype data and new query tools. Nucleic Acids Res. 2020 Jan 8;48(D1):D783–8.

2. Marnetto D, Huerta-Sánchez E. *Haplostrips*: revealing population structure through haplotype visualization. Methods Ecol Evol. 2017 Oct;8(10):1389–92.

3. Nemat-Gorgani N, Guethlein LA, Henn BM, Norberg SJ, Chiaroni J, Sikora M, et al. Diversity of KIR, HLA class I, and their interactions in seven populations of sub-Saharan Africans. J Immunol. 2019 May 1;202(9):2636–47.

4. Shkuri O, Israeli S, Tshuva Y, Maiers M, Louzoun Y. Efficient test for deviation from Hardy-Weinberg equilibrium with known or ambiguous typing in highly polymorphic loci. Brief Bioinform. 2024 Jul 25;25(5):bbae416.
